# Metagenomic resolution of spotted-fever group *Rickettsia tasmanensis* and novel DNA viruses in Australian wildlife ticks, with spatial modelling of *Rickettsia* exposure zones

**DOI:** 10.1101/2025.07.08.663691

**Authors:** Ernest J.M. Teo, Mary E. Petrone, Adam Stewart, Delaney Burnard, Stephen C. Barker, Rhys H Parry

## Abstract

Australia’s spotted fever group (SFG) rickettsiae cause significant illness, yet microbial diversity in wildlife ticks remains incompletely understood. Metagenomic sequencing of nine tick pools representing three Ixodidae species, *Ixodes tasmani*, *I. holocyclus*, and *Haemaphysalis bancrofti*, from Australian wildlife revealed a genome of a novel SFG Rickettsia species from *I. tasmani* ticks collected from koalas in New South Wales. Phylogenomic analysis confirmed this as a distinct species closely related to *Rickettsia tasmaniensis* fragments previously reported from Tasmanian devils. Additionally, we discovered four novel DNA Anellovirus species forming a new genus ‘Sintorquevirus’ and one Circovirus from likely vertebrate blood meals, revealing wildlife viral diversity. Species distribution modelling of *I. tasmani* for assessing *Rickettsia* risk revealed suitable habitat along Australia’s eastern coastlines, with overlap between vector distribution, marsupial hosts, and population centres. This identified potential Candidatus R. tasmanensis exposure zones in coastal regions where positive samples originated, highlighting areas for surveillance.

## Introduction

Ticks (Acari: Ixodida) are hematophagous ectoparasitic arthropods of mammals, birds, reptiles and amphibians. In Australia, 73 valid tick species have been described, belonging to seven of the eight subfamilies of tick species (Barker & Barker 2023), with 17 species capable of feeding on humans and domestic animals. The remaining 56 ticks mainly feed on birds, wild reptiles, and mammals (Barker & Barker 2023).

Within the *Ixodes* genus, *Ixodes holocyclus* causes tick paralysis in domestic animals, humans, and wildlife requiring bandicoots and other marsupials to sustain its life cycle (Barker and Walker, 2014; Barker & Barker 2023). *Ixodes tasmani* has the most widespread geographic distribution and the broadest range of hosts of any Australian tick, making it epidemiologically important for studies of tick-borne pathogens.

Ticks harbour complex microbiomes, including pathogenic bacteria transmitted via blood meals. In Australia, Q fever (*Coxiella burnetii*) and rickettsial infections (*Rickettsia* spp.) are the primary bacterial diseases transmitted by human-biting ticks. However, Australian ticks are also associated with complex clinical presentations that remain poorly understood. In 2018, the Australian National Health and Medical Research Council established a Targeted Call for Research into ‘Debilitating Symptom Complexes Attributed to Ticks’ (DSCATT), acknowledging that some Australians suffer chronic, debilitating symptoms following tick bites even when conventional diagnostics detect no specific pathogen. Additionally, ticks serve as effective xenosurveillance tools, enabling pathogen detection in wildlife hosts that are difficult to sample directly.

*Rickettsia* are obligate intracellular, Gram-negative bacteria transmitted vertically between invertebrates through life stages or transmitted horizontally between invertebrates and vertebrates during tick feeding. Phylogenetically, *Rickettsia* are categorised into four groups: the spotted fever group (SFG), the typhus group (TG), the ancestral group, and the transitional group, though clinically distinguished primarily as SFG or TG (Parola et al., 2013).

Four *Rickettsia* species cause human disease in Australia: *Rickettsia australis* (Queensland tick typhus), *Rickettsia honei* (Flinders Island spotted fever), *R. honei* subspecies *marmionii* (Australian spotted fever), and *Rickettsia typhi* (murine typhus) (Stewart et al., 2017b). *R. gravesii* has been identified in Australian ticks, though its pathogenicity requires further investigation (Graves and Stenos, 2009; Graves and Stenos, 2017). Despite substantial disease burden, particularly in tropical settings, no current disease surveillance systems monitor the incidence of *Rickettsia* spp. infection, and only Western Australia and Tasmania require notification (Stewart et al., 2017b; Stewart et al., 2019). Suboptimal diagnostic assays and a lack of consensus around definitions for disease also complicate accurate characterisation of disease epidemiology in Australia (Stewart and Stewart, 2021).

Risk factors for *Rickettsia* infection include occupational (farming, commercial harvesting) and recreational activities (bushwalking) (Stewart et al., 2017b). SFG *Rickettsia* infections typically occur within a 20-kilometer coastal band along eastern Australia, with recent studies documenting increasing incidence (Stewart et al., 2019): North Queensland identified 37 patients over 20 years, Southeast Queensland found 36 patients over 15 years (Stewart et al., 2017a), while Tasmania averaged 3 cases annually (0.6 per 100,000 population/year) over 6 years (Willis et al., 2019). The annual rate of SFG rickettsioses increased 8.5-fold from 2008 to 2012, reaching 14.3 cases per million (Drexler et al., 2016).

Species distribution modelling (SDM) provides a framework for projecting tick geographic ranges and assessing exposure risk by integrating tick occurrences with environmental variables, host distribution, and human population density. Metagenomic approaches can reveal novel microorganisms that conventional targeted molecular assays might miss (Barbosa et al., 2022). While previous studies have characterised viral and bacterial communities in Australian ticks using RNA sequencing (Chandra et al., 2021; Harvey et al., 2019; O’Brien et al., 2018), DNA-based metagenomic studies focusing on Australian wildlife ticks have been limited, making these approaches valuable for investigating emerging pathogens like novel *Rickettsia* species.

In this study, we employed high-throughput DNA metagenomic sequencing to characterise microbial communities in ticks collected from Australian wildlife. This enabled identification of a novel SFG *Rickettsia* species in *I. tasmani* ticks from koalas, and discovery of new viruses from *Anelloviridae* and *Circoviridae* families. We developed species distribution models for *I. tasmani*, integrating these with host occurrence data and human population centres to identify potential pathogen exposure risk zones. These combined genomic and spatial analyses expand our understanding of the diversity and geographic distribution of potential pathogens in Australian ticks, providing valuable insights for public health surveillance and risk management.

## Materials and Methods

### Tick collection, identification and genome sequencing

The data used in this study was generated from Burnard and Shao 2019, ticks were opportunistically collected by veterinarians and wildlife carers, stored in 70% ethanol and morphologically identified as previously described (Burnard et al., 2017). Tick specimens were collected from three sites, two in South-East Queensland and one in coastal New South Wales, spanning a geographic distance of approximately 800–1000 km. Nine tick pools consisting of three Ixodidae species, *I. tasmani* (n = 4), *I. holocyclus* (n = 4) and *H. bancrofti* (n = 1) (Table 1) were metagenome sequenced following previously described sample preparation methods (Taylor-Brown et al., 2016). The Australian Genome Research facility (AGRF) performed shotgun library preparation and sequencing on an Illumina HiSeq platform to produce between 25 and 32 million reads (Burnard and Shao, 2019).

**Table 1:** Metadata of tick samples used for high-throughput sequencing.

| ID | Tick species | Pool | Location | Host | Reads |
| --- | --- | --- | --- | --- | --- |
| A6T0 | *Ixodes tasmani* | 5 | South-East Queensland, Australia | Koala (*Phascolarctos cinereus*) | 30787648 |
| B6 | *Ixodes tasmani* | 5 | Coastal New South Wales, Australia | Koala (*Phascolarctos cinereus*) | 25720123 |
| C9 | *Ixodes tasmani* | 5 | Coastal New South Wales, Australia | Koala (*Phascolarctos cinereus*) | 30886771 |
| D3b | *Ixodes tasmani* | 1 | South-East Queensland, Australia | Long nosed bandicoot (*Perameles nasuta*) | 29226500 |
| B9b2 | *Ixodes holocyclus* | 1 | South-East Queensland, Australia | Brushtail possum (*Trichosurus vulpecula*) | 28211755 |
| E5b2 | *Ixodes holocyclus* | 1 | South-East Queensland, Australia | Spotted tail quoll (*Dasyurus maculatus*) | 32491495 |
| H4 | *Ixodes holocyclus* | 5 | South-East Queensland, Australia | Koala (*Phascolarctos cinereus*) | 26911342 |
| I7 | *Ixodes holocyclus* | 5 | South-East Queensland, Australia | Koala (*Phascolarctos cinereus*) | 26773592 |
| C9b2 | *Haemaphysalis bancrofti* | 1 | South-East Queensland, Australia | Eastern grey kangaroo (*Macropus giganteus*) | 27452720 |

### Metagenome assembly, contig binning and abundance quantification

Base-called FASTQ reads were trimmed for adapters and low-quality sequences using Trim Galore! (v0.6.7). The composition of each library was first analysed using CCMetagen (Marcelino et al., 2020), KMA (Clausen et al., 2018), using the NCBI nt database (June 2019), and then subject to *de novo* assembly using MEGAHIT (v1.2.9) (Li et al., 2015) and metaplasmidSPAdes (v3.15.4) to assemble circularised DNA contigs (Antipov et al., 2019). For detection of ribosomal RNA genes (5S, 16S and 23S), barrnap (v1.2.2) and BLASTn (Camacho et al., 2009) were utilised to identify mitochondrial, and bacterial contigs. Putative virus contigs were detected using BLASTx against a non-redundant local virus database previously described (Parry et al., 2021). For identification of *Rickettsia* contigs we utilised both MaxBin2 (v2.2.7) to cluster metagenomic contigs from the B6T sample into bins (Wu et al., 2016). The contigs from the putative *Rickettsia* bin was then further manually curated using BLASTn against the NCBI non-redundant (nr) database to exclude non-*Rickettsia* contigs.

For abundance estimates, reads were remapped to tick mitochondrial genomes (Burnard and Shao, 2019), mammalian host mitochondrial (*Macropus giganteus* Genbank: LK995454.1, *Phascolarctos cinereus* Genbank: NC_008133.1, *Trichosurus vulpecula* Genbank: CM021960.1, *Dasyurus hallucatus* Genbank: AY795973.1 *Perameles nasuta* Genbank: KJ868137.1) and pathogen contigs using Bowtie2 (v2.1.0) (Camacho et al., 2009).

### Virus genome annotation and phylogenetic analyses

Open reading frames of the novel DNA viruses identified here were predicted using the NCBI ORF finder (https://www.ncbi.nlm.nih.gov/orffinder/). Predicted protein sequences were analysed for protein domains using CD-Search against the Conserved Domain Database (Marchler-Bauer et al., 2011). For DNA folding of the circovirus stem loop, the Mfold (v4.7) web server was utilised under default settings (Zuker, 2003).

For phylogenetic placement of the novel ‘Sintorquevirus’ species and genus within the *Anelloviridae,* we downloaded 142 representative ORF1 protein coding sequences from 29 genera of anelloviruses, excluding *Gyrovirus* as their VP1 is not homologous to ORF1, and aligned using MAFFT (v7.475, L-INS-i algorithm) (Katoh et al., 2019). The ORF1 was then trimmed for ambiguous sequences using TrimAL (v 1.3) (Capella-Gutierrez et al., 2009), under the gappyout option. For the novel Circovirus we downloaded nucleotide sequences from 30 whole genomes and aligned the genomes using MAFFT (FFT-NS-I algorithm). IQ-TREE2 (v2.1.2) was then used to construct consensus maximum-likelihood phylogenetic trees from the alignments with substitution models using ModelFinder (Kalyaanamoorthy et al., 2017) using the Bayesian Information Criterion. Phylogenies were also built using the SH-aLRT test (-alrt 1000) and with 1000 ultrafast bootstraps (-B 1000). Resultant consensus trees were visualised using FigTree v1.4 (Andrew Rambaut, https://github.com/rambaut/figtree/).

### Identification of ‘Sintorquevirus’ reads in deposited koala high-throughput sequencing data

Koala (*Phascolarctos cinereus,* taxid: 38626) transcriptome libraries deposited on the SRA were queried using BLASTn from the six assembled anelloviruses. Libraries were considered anellovirus positive if they obtained greater than 10 reads with greater than 69% nucleotide identity. For visualisation of mapped RNA-Seq reads, the library with the greatest number of hits was downloaded and mapped to the anellovirus genomes using Bowtie2 with coverage calculated and visualised as previously described (Jansen van Vuren et al., 2021).

### *Candidatus* Rickettsia tasmanensis B6T genome annotation and phylogenetic analyses

Coding regions and noncoding RNAs (ncRNAs) of the assembled *Candidatus* Rickettsia tasmanensis B6T genome were annotated using the NCBI prokaryotic genome annotation pipeline (Tatusova et al., 2016). Genome completeness was assessed using BUSCO (v5.4.4) against the lineage dataset rickettsiales_odb10 (Creation date: 2020-03-06, number of genomes: 34, number of BUSCOs: 364) with the gene predictor prodigal used (Simao et al., 2015). CheckM analysis (v1.2.2) was also used to establish genome quality and was calculated on the Prokaryotic Genome Annotation Pipeline (PGAP) gene set with the *Rickettsia* CheckM marker set (Parks et al., 2015).

Whole genome genetic comparisons were analysed using OrthoANI as implemented in Orthologous Average Nucleotide Identity (OAT v0.9) (Lee et al., 2016) to measure the overall similarity between the *Rickettsia tasmaniensis* genome assembly and other Australian and global *Rickettsia* species. For *Rickettsia* multilocus sequence typing (MLST), we downloaded 16S rRNA/ssu, gltA, sca0/OmpA, sca5/OmpB and sca4 sequences from representative *Rickettsia* species and extracted sequences from *R. australis str_Phillips* (Dong et al., 2012), *R. gravesii* BWI-1 (Sentausa et al., 2013) and *R. honei* RB (Xin et al., 2012) genomes. The pairwise nucleotide similarity of a conserved 495bp fragment of ompB was calculated based on the imported MAFFT alignment in CLC Main Workbench (v 6.9.2).

### Species distribution modelling and risk assessment

To assess the potential geographical distribution of *Candidatus* R. tasmanensis and its risk for human exposure, we developed a species distribution model for *I. tasmani* using the species distribution model, MaxEnt (Phillips et al., 2017).

We assembled 942 occurrence records of *I. tasmani* from multiple sources: the Barker & Barker Collection; museum collections; Global Biodiversity Information Facility (GBIF, accessed March 2025); and published literature (Barker and Barker, 2023; Greay et al., 2016). Records with spatial accuracy of ≤10 km were retained (627 records). Records were then mapped onto climate data at a resolution of 0.05° × 0.05° (approximately 5.5 × 4.5 km at Sydney), and filtered to remove pixel duplicates (283 locations). These were spatially thinned at 8km using spThin (Aiello-Lammens et al., 2015) resulting in 233 final locations for modelling.

We selected seven bioclimatic variables based on their relevance to tick ecology: mean annual temperature (bio01), mean diurnal range (bio02), temperature seasonality (bio04), annual precipitation (bio12), precipitation seasonality (bio15), precipitation of driest quarter (bio17), and mean vapor pressure deficit (vpd). These variables are also commonly used in tick distribution modelling (Estrada-Peña et al., 2008; Hahn et al., 2016; Leta et al., 2013; Ma et al., 2024; Ma et al., 2023; Pascoe et al., 2022; Perret et al., 2000; Raghavan et al., 2019; Randolph and Storey, 1999; Rochlin et al., 2023; Teo et al., 2024a; Teo et al., 2021a; Teo et al., 2023; Teo et al., 2024b; Teo et al., 2021b; Xu et al., 2024; Zemtsova et al., 2016). Climate data from 1991-2020 were obtained from SILO (https://www.longpaddock.qld.gov.au/silo/) (Jeffrey et al., 2001), and the bioclimatic variables were prepared using the predicts package (Hijmans et al., 2024).

### Model calibration and evaluation

For MaxEnt parameter optimisation, we used ENMeval (Kass et al., 2021) to test combinations of feature classes (linear, quadratic, and hinge) (Phillips et al., 2017) and regularisation multipliers (1 to 5 at 0.5 intervals). We generated 10,000 random background points within a 100 km buffer around the occurrence points. For cross-validation, the data were split into five spatial blocks using blockCV (Valavi et al., 2019). Models were selected using the continuous Boyce Index (scale from -1 to 1), which measures agreement between model predictions and test data (Hirzel et al., 2006; Low et al., 2021). The 10th percentile training presence threshold was applied to differentiate suitable from unsuitable habitats.

### Risk area identification

To identify potential areas of zoonotic transmission risk, we overlaid three datasets: vector distribution (derived from our MaxEnt model), host distribution, and human population density. Occurrence data for all 48 marsupial host species of *I. tasmani* (Barker and Barker, 2023) were retrieved from the Atlas of Living Australia (March 6, 2025) using the galah package (Westgate et al., 2025). Only records from 1991 onward were retained to match the temporal extent of climate and human population data.

For relative host abundance estimation, we removed duplicate records and summed host occurrences at 0.05° × 0.05° resolution. We then downloaded occurrence records for all relevant marsupial families (Acrobatidae, Burramyidae, Dasyuridae, Macropodidae, Peramelidae, Petauridae, Phalangeridae, Phascolarctidae, Potoroidae, Pseudocheiridae, and Vombatidae) to create a ‘target-group’ background (Barber et al., 2022). A list of marsupial hosts and the number of records retrieved are provided in the Supplementary File 1. Relative abundance was calculated by dividing the host count by the target-group count in each grid cell.

Human population data were obtained from the Australian Bureau of Statistics (2023) at 1km² resolution, then resampled to the EPSG:4326 projection. City and administrative region boundaries followed the Australian Bureau of Statistics’ “Greater Capital City Statistical Areas” and “Statistical Areas Level 4” (Australian Bureau of Statistics, 2021).

## Results

### Overview of tick collection and metagenomic sequencing

Nine tick pools from three Ixodidae species: *Ixodes tasmani* (n = 4), *Ixodes holocyclus* (n = 4), and *Haemaphysalis bancrofti* (n = 1) were analysed (Table 1). Ticks were collected from native Australian marsupials, including koalas (*Phascolarctos cinereus*), possums (*Trichosurus vulpecula*), bandicoots (*Perameles nasuta*), quolls (*Dasyurus maculatus*), and kangaroos (*Macropus giganteus*). Sequencing yielded between 25.7 and 32.5 million reads per sample (mean: 28.7 million reads).

Reads were taxonomically classified using CCMetagen (Fig. 1). Most assigned reads were eukaryotic (50-98%), with arthropod DNA constituting between 25-75% of eukaryotic reads. Marsupial DNA typically matched the host species from which ticks were collected, with one exception: sample E5b2 (*I. holocyclus* from spotted-tailed quoll), was misassigned to wombat, likely due to limited *Dasyurus* genomic representation in the NCBI database. Three samples (D3b, B9b2, C9b2) contained no marsupial reads, potentially reflecting recent host attachment. No human reads were detected. Our analysis revealed several bacterial taxa, including the tick endosymbiont *Midichloria* in *I. holocyclus* samples, *Lonepinella* in one *I. holocyclus* sample, and *Rickettsia* in *I. tasmani* from koala.

**Figure 1.**
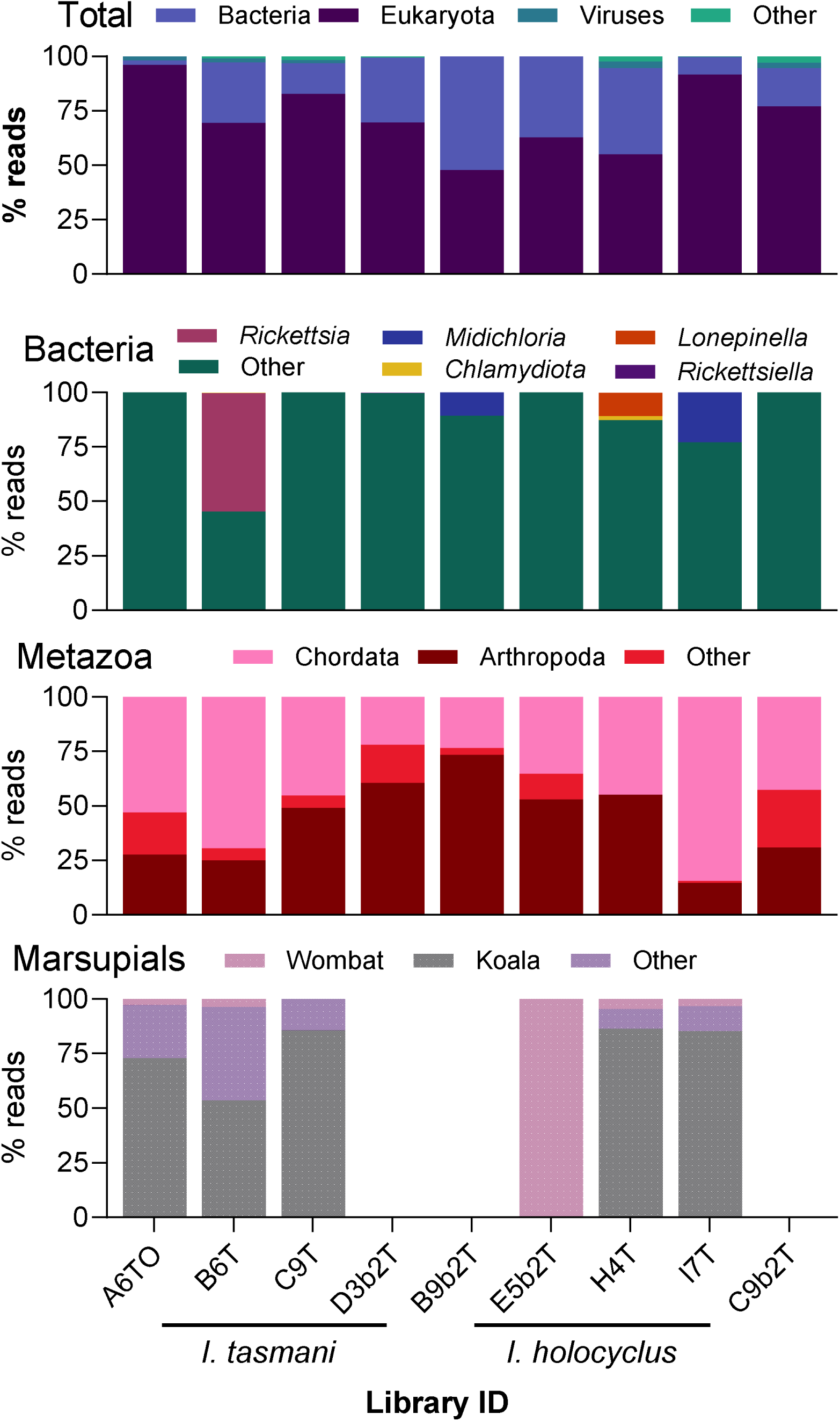
Taxonomic composition of metagenomic libraries from Australian ticks. Reads from each library were assigned to major taxonomic categories (Bacteria, Eukaryota, Viruses and other) using CCMetagen NCBI (nt_no_env_11jun2019) database, top panel. The relative abundance of Bacteria and Metazoa, along with the subcategory of Marsupial within Metazoa are shown as percentage of assigned reads per category. Sample IDs are shown on the x-axis.

### Assembly of a novel *Rickettsia* genome species from *I. tasmani*

Metagenomic analysis and binning of contigs from sample B6T enabled the assembly of a draft genome for a previously uncharacterized *Rickettsia* species isolated from *I. tasmani* ticks collected from koalas. Quality assessment revealed substantial genome recovery, with a CheckM score of 97.15% and a BUSCO analysis indicating 89.8% completeness based on the Rickettsiales dataset. The assembly comprises 97 scaffolds, totalling 1,331,613 bp with a GC content of 32.49%. While demonstrating moderate contiguity, as reflected by an N50 of 46,430 bp and an L50 of 9 scaffolds, the genome quality metrics are comparable to those of other Australian *Rickettsia* species (Table 2). Annotation via the NCBI Prokaryotic Genome Annotation Pipeline revealed 1,654 total genes, including 1,434 protein-coding genes, 42 RNA genes (4 rRNAs, 34 tRNAs, 4 ncRNAs), and 178 pseudogenes.

**Table 2:** Genomic features of Australian *Rickettsia* genomes. BUSCO Score abbreviations: C = Complete, S = Complete and single-copy, D = Complete and duplicated, F = Fragmented, M = Missing.

| Feature | *R. tasmaniensis* B6T<br>GCA_036857515 | *R. australis* Phillips<br>GCF_000273745 | *R. gravesii* BWI-1<br>GCF_000485845 | *R. honei* RB<br>GCF_000263055 |
| --- | --- | --- | --- | --- |
| CheckM Score | 97.15% | 95.6% | 99.05% | 99.31% |
| BUSCO Score | C:89.8%<br>[S:89.8%,D:0.0%]<br>F:1.6%,M:8.5% | C:89.3%<br>[S:89.3%,D:0%]<br>F:1.4%,M:9.3% | C:92.0%<br>[S:92.0%,D:0%]<br>F:0.5%,M:7.4% | C:92.6%<br>[S:92.6%,D:0%],<br>F:0.0%,M:7.4% |
| Genome Size | 1.33 Mb | 1.32 Mb | 1.34 Mb | 1.26 Mb |
| GC Content | 32.49% | 32.26% | 32.23% | 32.42% |
| Scaffolds | 97 | 59 | 29 | 11 |
| N50 | 46,430 bp | 57,972 bp | 71,065 bp | 163,849 bp |
| L50 | 9 | 9 | 8 | 3 |
| Total Genes | 1,654 | 1,478 | 1,575 | 1,494 |
| Protein-coding Genes | 1,434 | 1,267 | 1,364 | 1,309 |
| RNA Genes | 42 | 40 | 41 | 40 |
| rRNAs (5S, 16S, 23S) | 1, 2, 1 | 1, 1, 1 | 1, 1, 1 | 1, 1, 1 |
| tRNAs | 34 | 33 | 34 | 33 |
| ncRNAs | 4 | 4 | 4 | 4 |

Phylogenomic analysis revealed a distinct lineage within the *Rickettsia* spotted fever group (Fig. 2A). Average nucleotide identity (ANI) analysis (Supplementary Fig. 1) showed <99.19% OrthoANI similarity with known *Rickettsia* species, confirming novel species status with established thresholds for species demarcation (Diop et al., 2020). The closest relatives based on ANI values were *Rickettsia slovaca str DCWPP* (97.7%) and *Rickettsia africae ESF5* (97.40%).

**Figure 2.**
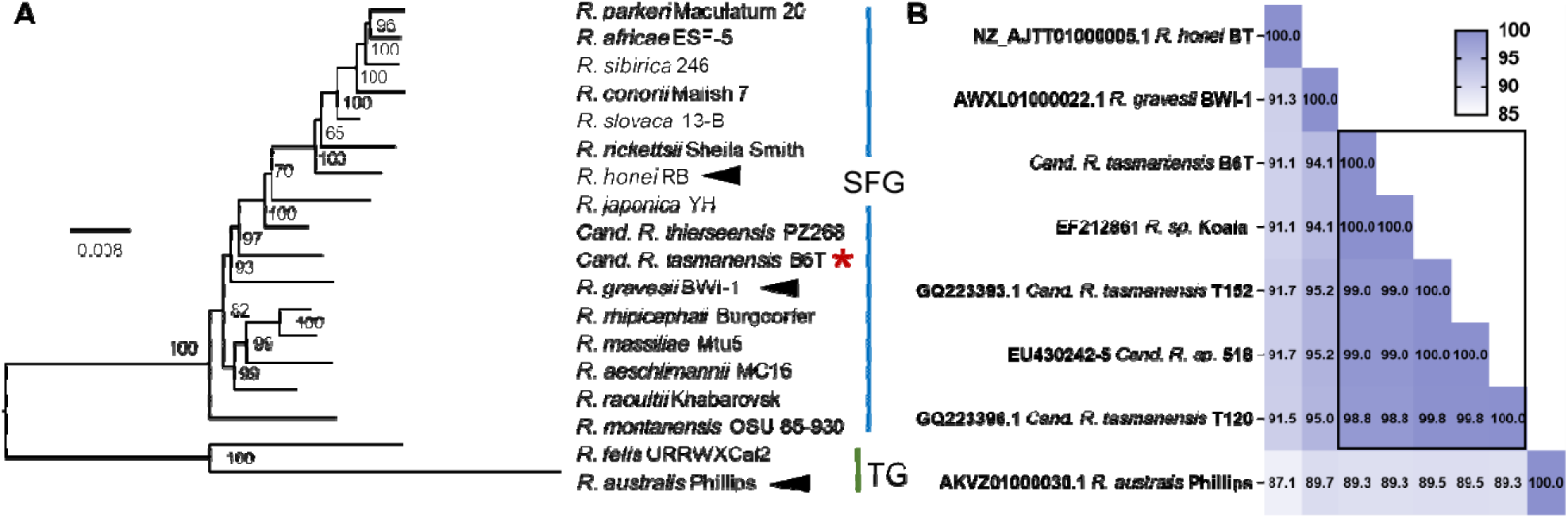
Phylogenomic analysis and sequence similarity of ompB of *Candidatus* Rickettsia tasmanensis. A) Phylogenetic tree based on multilocus sequence typing (MLST) analysis of rickettsial species using 16S rRNA, gltA, ompA, ompB, and sca4 genes, showing the phylogenetic placement of *Candidatus* R. tasmanensis within the spotted fever group (SFG). Branch length corresponds to nucleotide substitutions per site, model used was GTR+F+R2. B) Heatmap of all Australian SFG species and PCR fragments showing pairwise nucleotide similarity of a conserved 495bp fragment of the ompB gene. Samples from *I. tasmani* and likely to be *R. tasmaniensis* are indicated with a box.

### Harmonizing previous genetic studies on *Rickettsia* species in Australia

Previous studies have identified *Rickettsia* gene fragments from I. tasmani in both mainland Australia and Tasmania. Our ompB sequence from our assembly showed 100% identity to sequences (GenbankID: EF212861) from *I. tasmani* ticks collected from koalas in Port Macquarie (Vilcins et al., 2008) and 98.8-99.0% similarity to *Candidatus* R. tasmanensis from Tasmanian devils, *Rickettsia* sp. 518 (GenbankID: EU430242), *Candidatus* R. tasmanensis T120 (GQ223396), *Candidatus* R. tasmanensis T152-E (GenbankID: GQ223393) (Izzard et al., 2009)(Vilcins et al., 2009). However, only gene fragments for this putative species had been characterised until now. Our genome assembly confirms these previous detections represent a single, distinct species, formally establishing it as the fourth SFG *Rickettsia* species on mainland Australia.

### Exploring geographic distribution and potential risk areas for *Candidatus R. tasmanensis*

To assess the potential public health significance of *Candidatus* R. tasmanensis, we developed species distribution models for its vector *I. tasmani,* to identify geographic regions where human exposure might occur. Our species distribution model for *I. tasmani* demonstrated high predictive performance (Boyce Index: 0.93, AUC: 0.79), across five cross-validation sets, indicating strong agreement between model projections and test data.

The MaxEnt model revealed high climatic suitability for *I. tasmani* along the eastern and southeastern coasts of mainland Australia and throughout Tasmania (Fig. 3A). Other suitable regions included the Fleurieu Peninsula and Kangaroo Island in South Australia, as well as portions of southern Western Australia. Notably, the projected suitable range encompasses Australia’s three most populous cities: Brisbane, Sydney, and Melbourne, as well as Hobart, Tasmania.

**Figure 3.**
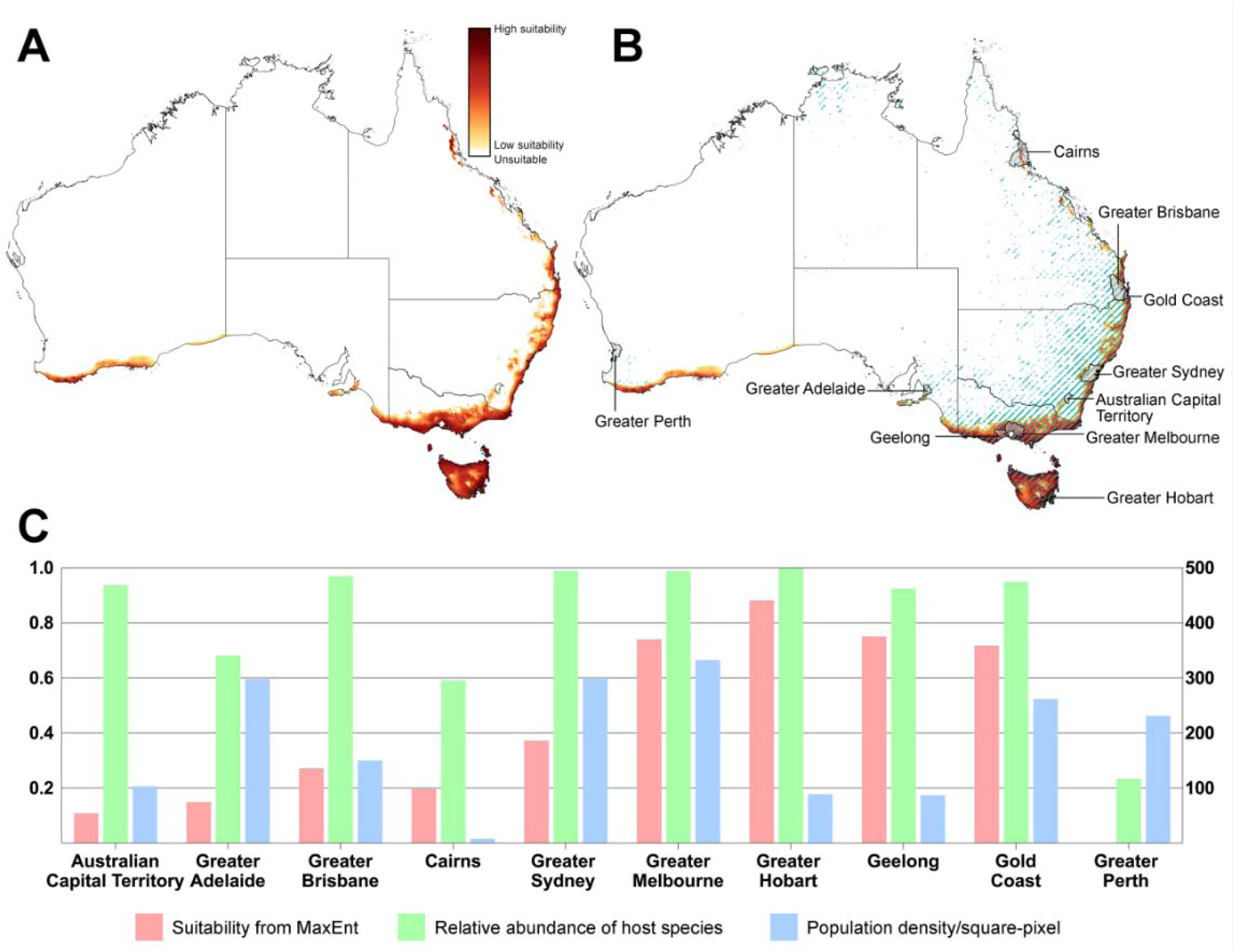
Geographic distribution and risk assessment for *Candidatus* R. tasmanensis in Australia. A) Potential geographic range of *Ixodes tasmani* projected from the MaxEnt model. White indicates unsuitable habitat, with darker colors indicating increasing climatic suitability. B) Distribution of *I. tasmani* (gradient colors) overlaid with marsupial host occurrence (blue lines), highlighting areas of potential transmission. Major population centres are labelled C) Comparison of climatic suitability from MaxEnt (red), relative abundance of host species (green), and human population density (blue) across major Australian cities and regions, identifying potential public health risk zones.

Overlaying vector distribution with host marsupial occurrence revealed substantial overlap in coastal New South Wales and Queensland (Fig. 3B). These zones coincide with areas of moderate to high human population density, creating potential interfaces for zoono transmission. However, these analyses estimate potential interaction zones based on climatic suitability and host occurrence rather than predicting absolute transmission risk, as actual disease transmission depends on additional factors including local infection prevalence, host competence, and human behaviour patterns which are beyond the scope of this model (Elith and Leathwick, 2009).

City-level analysis revealed that Greater Hobart had the highest climatic suitability (0.883) and host abundance (0.999) for *I. tasmani*, despite having a lower human population density than mainland capital cities. Greater Melbourne had high climatic suitability (0.740), and the highest human population density (333 persons per square pixel), along with substantial host abundance (0.988). Relative abundance represents suitable marsupial hosts relative to their taxonomic families, not actual host numbers.

This convergence of suitable tick habitat, marsupial hosts, and human populations creates potential exposure “hotspots” for *Candidatus* R. tasmanensis, particularly in coastal eastern Australia. The Port Macquarie region, where our *Rickettsia*-positive samples were collected, falls within these high-risk zones, validating our findings.

### A novel anellovirus genus ‘Sintorquevirus’ infects Australian marsupials

Anelloviruses (family *Anelloviridae*), are circular, negative single-stranded DNA viruses with genomes range in size from 1.6-3.9 kb in length (Varsani et al., 2021). Anelloviruses are thought to have a worldwide distribution in vertebrates. Anelloviruses have also been found in blood-feeding arthropods (Ng et al., 2011) including ixodid ticks (Waits et al., 2018) but are likely derived from the vertebrate blood meal of their host. In Australia A*nelloviridae*-like transcripts have been previously reported in Tasmanian devil samples (Harding et al., 2021).

We assembled anellovirus contigs from six of our eight samples (Fig. 4A), of which five were complete and circularised, and one was linear and not supported by circularisation. BLASTx analysis of putative anelloviruses suggested multiple but divergent hits with the closest relatives being the Torque teno tupaia virus (bit score: 97.1, Query cover: 36%, E-value: 1e-16, identity: 25.41%, GenbankID: YP_009505746), which was discovered in sera from northern treeshrews (*Tupaia belangeri*) in Yunnan Province, China (Okamoto et al., 2001) and Giant panda anellovirus (bit-score: 93.2, Query cover: 19%, E-value: 3e-15, identity: 28.50%, GenbankID: QZE11970) (Zhao et al., 2022) from Sichuan Province, China. All encoded a large ORF1 containing the viral capsid protein domain (accession: pfam02956, E-value range between 3.7E-19 and 8.8E-10) at the N-terminal region of ORF1 (Fig. 4A).

**Figure 4.**
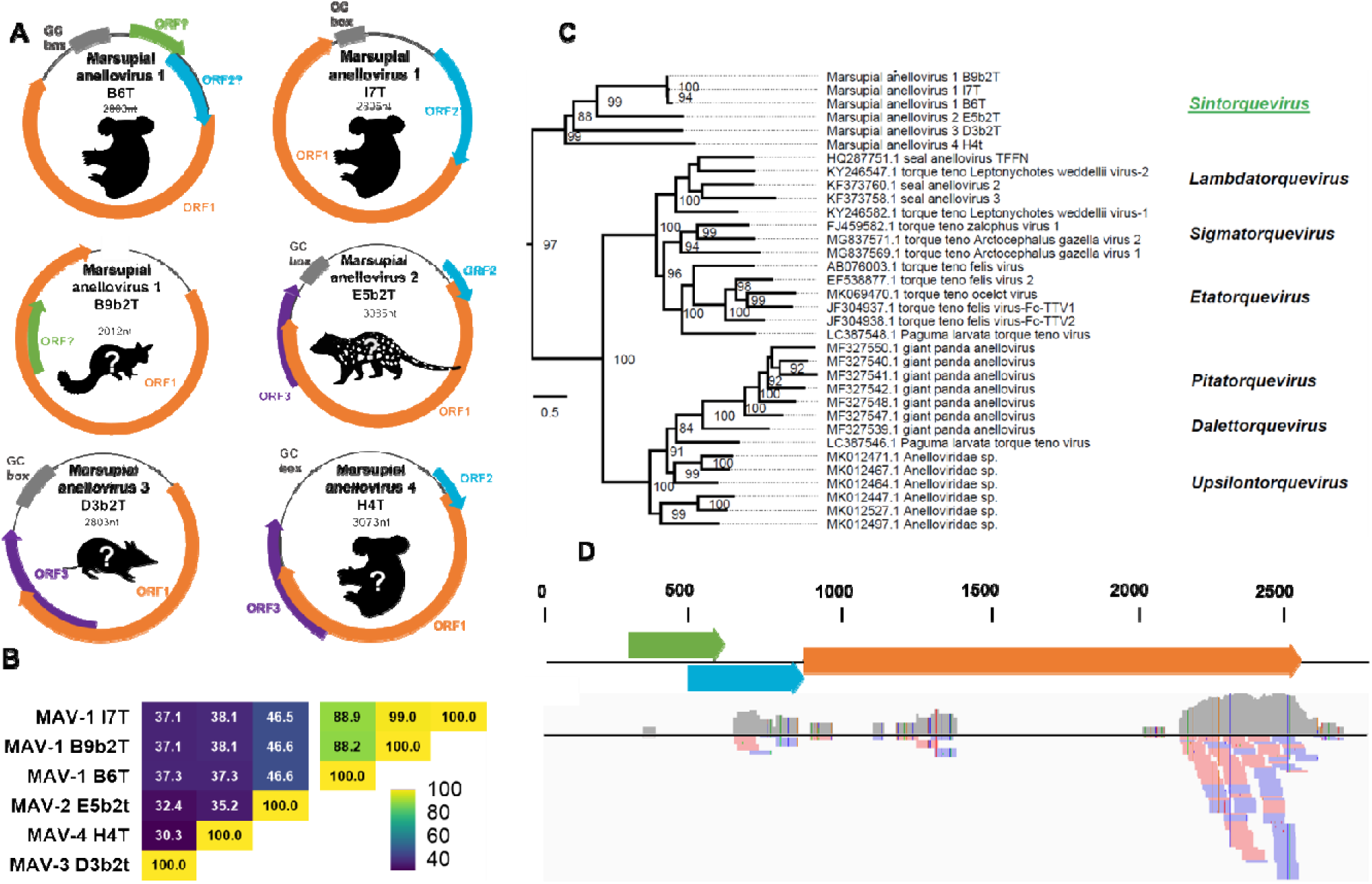
A novel genus of anelloviruses, ‘Sintorquevirus’ is present in Australian marsupials. A) Predicted genomic organisation of newly discovered anelloviruses. B) Pairwise nucleotide similarity between the ORF1 of anelloviruses and strains C) Truncated consensus maximum likelihood phylogenetic placement of the four novel marsupial anelloviruses (MAV) in the ‘Sintorquevirus’ genus, full tree available in Fig. S2. Branch length corresponds to nucleotide changes per position, model selection was GTR+I+G4. D) The host status of Marsupial anellovirus 1 B6T strain in Koalas is

*Anelloviridae* comprises 30 genera (Varsani et al., 2021), with species demarcation requiring <69% ORF1 nucleotide identity. All six assembled anellovirus ORF1 lacked BLASTn similarity to any published anellovirus sequences, indicating novel species. Pairwise alignment revealed three contigs (IT3, B9b2T, B6T) shared 88.2-99.0% identity (Fig. 4B) representing one species, while the remaining three were unique, yielding four novel species total. Maximum-likelihood phylogenetic analysis of ORF1 sequence with representatives from all 30 genera of *Anelloviridae* (Fig. 4C) showed these anelloviruses clustered in a divergent clade (Fig. 4C, Supplementary Fig. 2), indicating they belong to a completely different genus.

*Anelloviridae* nomenclature uses Greek and Phoenician alphabet prefixes with “torquevirus” as suffix. Using the Phoenician letter šīn, we define the new genus ‘Sintorquevirus’. Species epithets comprise the first five letters of the host family name plus a number. We propose four species: *Sintorquevirus ixodi1*, *S. ixodi2*, *S. ixodi3*, and *S. ixodi4*, with virus names Marsupial anellovirus 1-4.

To confirm marsupial association rather than tick contamination, we screened 182 koala RNA-Seq datasets from the Sequence Read Archive. Six datasets (3.2%) contained reads with 90-100% identity to marsupial anelloviruses. One koala PBMC sample (SRR10337972) with >200 reads was mapped to marsupial anellovirus 1, revealing transcriptional activity (Fig. 4D). This confirms these anelloviruses are likely associated with the blood meals when taken from Australian marsupials, and unlikely to represent bona fide tick viruses.

### A novel circovirus identified from an *I. tasmani* tick infesting a long-nosed bandicoot

*Circoviridae* are vertebrate viruses with circular, single-stranded DNA genomes (1.7-2.1 kb). Circovirus genomes typically encode two main genes: the replicase (rep), and the capsid (cp) gene, which encodes the structural proteins that form the virus capsid (Breitbart et al., 2017).

We discovered a 2050nt circular contig in the *I. tasmani* sample from the long-nosed bandicoot (*Perameles nasuta*). BLASTn analysis of the contig revealed closest similarity to Canary circovirus (79.03% identity,GenbankID AJ301633.2) from the Atlantic canary (*Serinus canaria*) (Todd et al., 2001). Since *Circoviridae* species demarcation requires ≥80% genome-wide nucleotide sequence identity, this circovirus contig likely represents a novel virus, which we have named Hard Tick Associated Cyclovirus D3b (HTC-D3b).

Genome annotation revealed typical ambisense organization with viral replicase (rep) gene (Accession: cl03577, Interval: 9-93, E-value: 5.80E-20) and RNA helicase (Accession: pfam00910, Interval: 161-244, E-value: 3.00E-18) on the virion strand, and capsid gene (accession: pfam02443, interval: 41-243, E-value: 6.95E-109) on the complementary strand (Fig. 5A). The genome contained conserved origin of replication elements including a stem-loop structure with the nonanucleotide motif [(T/n)A(G/t)TATTAC] (Fig. 5B).The genome contained conserved origin of replication elements including a stem-loop structure with the nonanucleotide motif [(T/n)A(G/t)TATTAC] (Fig. 5B) (Wang et al., 2018).

**Figure 5.**
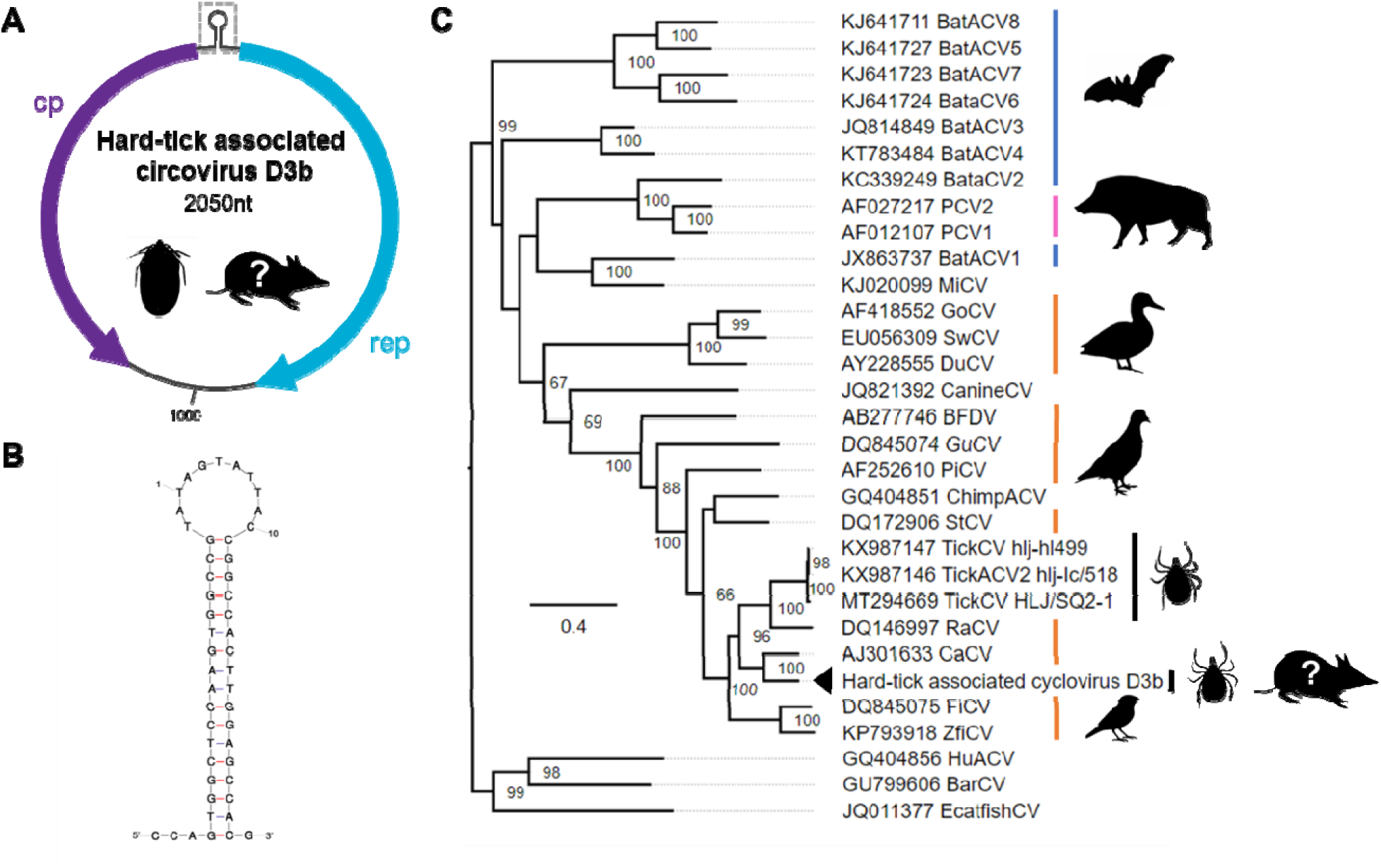
A novel cyclovirus identified in the blood meal of *I. tasmani*. A Genome schematic showing the capsid (cp, purple) and replicase (rep, blue). Stem loop structure of the ori C. Consen us indicates bat hosts, pink indicates porcine and black indicates tick hosts. Branch length corresponds to the number of amino acid changes per position; the protein substitution model is LG+R7.

### Detection of Trypanosoma gilletti

In one of the samples from koalas, we detected *Trypanosoma gilletti*. BLASTn analysis revealed 100% identity with *T. gilletti* (GenbankID: GU966589.1) previously documented in *Phascolarctos cinereus* (McInnes et al., 2011). In our assembly, we also found abundant kinetoplast DNA (kDNA), the mitochondrial genome of trypanosomatids. kDNA consists of a few dozen maxicircles and several thousand minicircles, all catenated topologically to form a two-dimensional DNA network. Minicircles are heterogeneous in size and sequence among species. They present one or several conserved regions that contain three highly conserved sequence blocks: CSB-1 (10 bp sequence), CSB-2 (8 bp sequence), and CSB-3 (12 bp sequence, also known as the Universal Minicircle Sequence).

We assembled 30 variants of the *T. gilletti* minicircle, all of which contained the characteristic conserved sequence blocks (Table S1). This finding confirms the presence of *T. gilletti* in koalas and suggests that *I. tasmani* ticks may play a role in the transmission of this parasite between koala populations.

## Discussion

We identified novel microorganisms in Australian ticks using high-throughput DNA sequencing, with the most significant finding being the first complete genome (BUSCO Score: 89.8%) of a novel spotted fever group *Rickettsia* species, *Candidatus* R. tasmanensis, from *I. tasmani* ticks collected from koalas. Phylogenomic analysis revealed an average nucleotide identity of less than 99.19% with known *Rickettsia* species, confirming this as a distinct species. This genomic characterisation extends previous molecular studies that had only identified gene fragments (Izzard et al., 2009; Vilcins et al., 2009; Vilcins et al., 2008) establishing it as the fourth SFG *Rickettsia* species on mainland Australia.

Our species distribution modeling demonstrates substantial geographic overlap between *I. tasmani* distributions, key marsupial hosts, and human population centres, particularly along the eastern Australian coastline where SFG *Rickettsia* infections typically occur (Stewart et al., 2017b). Our modelling identified several high-risk regions where environmental suitability, high marsupial host abundance, and significant human populations converge. Greater Hobart demonstrated the highest suitability and host abundance metrics, while Melbourne’s combination of substantial tick suitability, host abundance, and high human population density creates conditions for potential zoonotic exposure. High-risk areas in coastal New South Wales, overlapping with where our *Rickettsia*-positive samples were collected, should be prioritized for surveillance programs.

The ecological flexibility and broad host range of I. tasmani significantly increases zoonotic transmission risk. Our analysis revealed 48 marsupial species associated with this tick, allowing it to interface between wildlife, domestic animals, and humans in populated coastal regions. Previous studies documented high SFG *Rickettsia* prevalence in domestic animals near Launceston, Tasmania, with 10 of 16 tick samples testing positive (Izzard et al. 2009), further highlighting the potential for transmission.

Climate change may alter these *potential* risk patterns by shifting climatic suitability zones, potentially expanding the geographic range of *I. tasmani* and *Candidatus* R. tasmanensis into new areas (Robinson et al., 2015). However, predicting future *actual* risk remains complex, depending on concurrent changes in host communities, land use, and pathogen adaptation (Beard et al., 2016). This underscores the importance of continued monitoring and adaptive public health strategies across eastern Australia.

Current diagnostic approaches for *Rickettsia* infections include serological methods and nucleic acid amplification testing, though both have timing limitations: antibody development doesn’t occur until the second week of illness, while *Rickettsia* DNA in blood is short-lived and occurs early during illness (Stewart and Stewart, 2021). Advanced molecular methods, including high-throughput sequencing for clinical metagenomics, offer potential for improved detection (Graham et al., 2017). *Candidatus* R. tasmanensis is likely to respond to standard treatment protocols, typically doxycycline, with a seven-day course showing rapid response (24-48 hours) and complete tick removal as a critical management step.

The discovery of multiple viral species underscores the complexity of tick-associated microbiomes and the value of xenosurveillance for wildlife pathogen surveillance. While previous RNA metatranscriptomics studies detected RNA viruses, our DNA-based approach revealed viruses that are unlikely to be detected through RNA metatranscriptomics, which are likely associated with vertebrate blood meals rather than active tick infections. We report four novel ‘Sintorquevirus’ species within the *Anelloviridae* family and a unique circovirus, expanding known viral diversity in Australian wildlife through tick xenosurveillance. Verification of viral presence through RNA-seq data, particularly for marsupial anellovirus in koala samples, demonstrates the importance of orthologous analysis approaches.

This study expands understanding of tick-associated microorganisms in Australia. The findings highlight the dynamic nature of vector-borne disease ecology, emphasising the need for continued surveillance and molecular approaches to understand potential public health risks. Future research should focus on determining the pathogenic potential of the assembled *Rickettsia* species and exploring the ecological relationships between ticks, their hosts, and associated microorganisms.

Dr. Parry is a research fellow at the School of Chemistry and Molecular Biosciences, University of Queensland, Australia. His research topics cover virus bioinformatics and molecular characterisation of metagenomics-identified viruses.

## Ethics approval for sampling

Tick collection was carried out opportunistically on animals presenting for care or collected as part of other animal health investigations. The approval to complete tick sampling and *Chlamydia* screening was sought and approved by the University of the Sunshine Coast Animal Ethics Committee (ANS1539).

## Data Availability

Raw sequencing data produced for this study have been deposited in the Sequence Read Archive (SRA) in the NCBI database under the BioProject accession number PRJNA936717. The draft genome sequence for *Candidatus* Rickettsia tasmanensis strain:B6T is available under the BioProject accession PRJNA914631 and accession number GCA_036857515.

## Supplementary Files

Supplementary File 1 A list of marsupial hosts and the number of records retrieved for the analysis.

## Author contributions

R.H.P. and D.B. conceived the study. D.B. provided the metagenomic data generated from Burnard and Shao, 2019. D.B. obtained funding. R.H.P, E.T. and M.E.P. performed the analyses. R.H.P, M.E.P and E.T. provided visualisation. R.P. drafted the manuscript. S.C.B. provided resources. A.S., provided validation and technical interpretation. All authors reviewed and edited the manuscript.

## Acknowledgements

This research utilized data generated by Burnard and Shao 2019, which was funded by the Holsworth Wildlife Research Endowment – Equity Trustees Charitable Foundation & the Ecological Society of Australia, awarded to DB. The authors wish to acknowledge Arvind Varsani’s assistance in *Anellovirus* nomenclature. This work utilised the Australian Galaxy service (https://usegalaxy.org.au/). We acknowledge the technical assistance and proofing suggestions of John Stenos, Stephen Graves and Tarka Raj Bhatta from the Australian Rickettsial Reference Laboratory. RHP is funded by a Australian Research Council DECRA Fellowship (DE260101042).

---

**Supplementary Figure 1.**
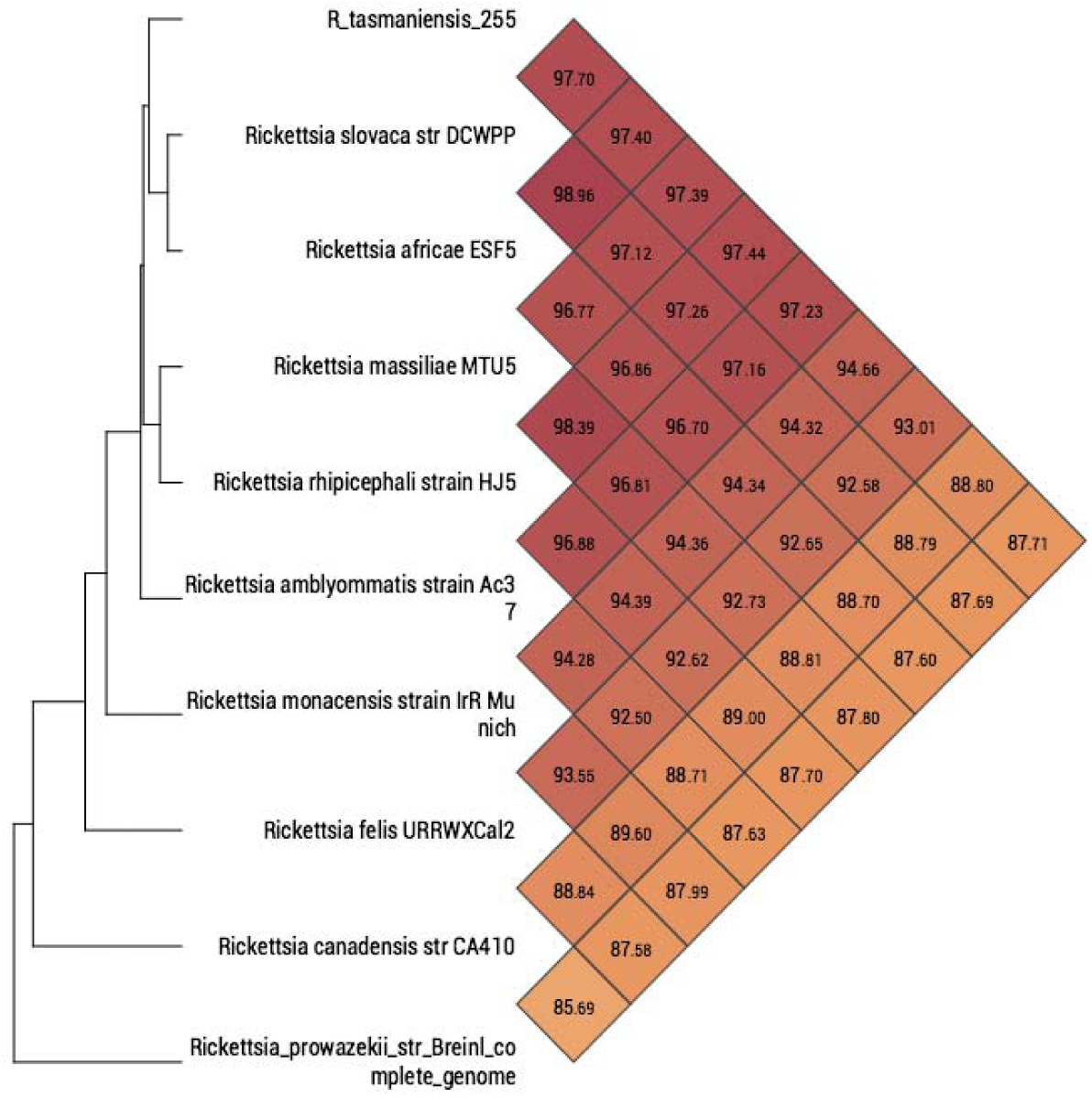
Phylogenetic tree and whole-genome Average Nucleotide Identity (ANI) heatmap of *Candidatus R. tasmanensis* and related *Rickettsia* species. Phylogenetic relationships are based on whole-genome sequence alignment, with accompanying ANI values demonstrating the genetic distinctiveness of *Candidatus R. tasmanensis* from other Spotted Fever Group *Rickettsia* species, with ANI values ranging from 85-98%.

**Supplementary Figure 2.**
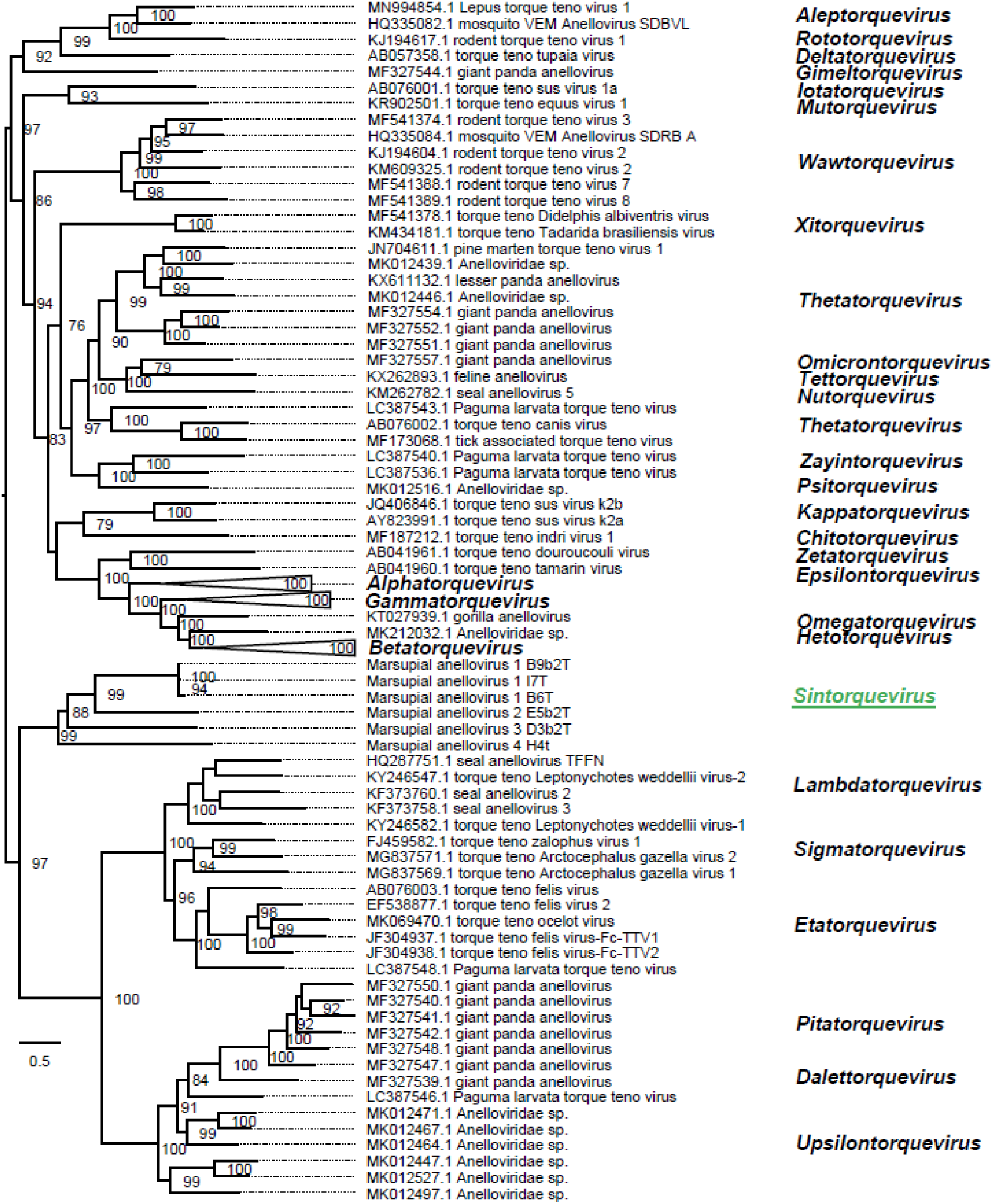
Maximum likelihood phylogenetic tree of the *Anelloviridae* family based on ORF1 protein sequences. The proposed ‘Sintorquevirus’ genus, representing novel marsupial anelloviruses, is highlighted in green. Bootstrap support values above 75 are shown. Branch length corresponds to amino acid changes per position.

**Supplementary Table 1.** Comparative analysis of *Trypanosoma* minicircle conserved sequence blocks (CSB). D1-2: average distance (in bp) between CSB-1 and CSB-2; D2-3: average distance (in bp) between CSB-2 and CSB-3 or UMS.

| Organism | CSB-1 | D1-2 | CSB-2 | D2-3 | CSB-3 or UMS | UMS per minicircle | Sequence reference |
| --- | --- | --- | --- | --- | --- | --- | --- |
| *T. gilletti* | AGGGGCGTTC | 30 | CCCCGTAT | 48 | GGGGTTGGTGTA | 2,3,4,5 | **This study** |
| *T. cruzi* Y strain | AGGGGCGTTC | 30 | CCCCGTAC | 47 | GGGGTTGGTGTA | 4 | TCU07845 |
| *T. copemani* G1 | AGGGGCGTGC | 28 | CCCCGTAC | 46 | GGGGTTGGTGTA | 2, 4 | MG948556<br>MG948555 |
| *T. lewisi* | GAGGGCGTTC | 32 | CCCCGTAT | 46 | GGGGTTGGTGTA | 2 | M17996 |
| *T. congolense* | AAGGGCGTTC | 29 | TCCCGTAC | 47 | GGGGTTGGTGTA | 1 | Nasir et al.<br>(1987) |
| *T. equiperdum* | ATGGGCGTGC | 21 | TCACGTGC | 38 | GGGGTTGGTGTA | 1 | Barrois et al.<br>(1981) |
| *T. brucei* | ATGGGCGTGC | 20 | TCCCGTGC | 41 | GGGGTTGGTGTA | 1 | Jasmer and<br>Stuart (1986) |

## References

Aiello-Lammens, M.E., Boria, R.A., Radosavljevic, A., Vilela, B., Anderson, R.P., 2015. spThin: an R package for spatial thinning of species occurrence records for use in ecological niche models. Ecography 38, 541–545.

Antipov, D., Raiko, M., Lapidus, A., Pevzner, P.A., 2019. Plasmid detection and assembly in genomic and metagenomic data sets. Genome Research 29, 961–968.

Barber, R.A., Ball, S.G., Morris, R.K.A., Gilbert, F., 2022. Target-group backgrounds prove effective at correcting sampling bias in Maxent models. Diversity and Distributions 28, 128–141.

Barbosa, A.D., Long, M., Lee, W., Austen, J.M., Cunneen, M., Ratchford, A., Burns, B., Kumarasinghe, P., Ben-Othman, R., Kollmann, T.R., Stewart, C.R., Beaman, M., Parry, R., Hall, R., Tabor, A., O’Donovan, J., Faddy, H.M., Collins, M., Cheng, A.C., Stenos, J., Graves, S., Oskam, C.L., Ryan, U.M., Irwin, P.J., 2022. The Troublesome Ticks Research Protocol: Developing a Comprehensive, Multidiscipline Research Plan for Investigating Human Tick-Associated Disease in Australia. Pathogens 11, 1208.

Barker, S.C., Barker, D., 2023. Ticks of Australasia: 125 species of ticks in and around Australia. Zootaxa 5253, 1–670.

Barker, S.C., Walker, A.R., 2014. Ticks of Australia. The species that infest domestic animals and humans. Zootaxa 3816, 1–144.

Beard, C.B., Eisen, R.J., Barker, C.M., Garofalo, J.F., Hahn, M., Hayden, M., Monaghan, A.J., Ogden, N.H., Schramm, P.J., 2016. Ch. 5: Vectorborne Diseases, The Impacts of Climate Change on Human Health in the United States: A Scientific Assessment. U.S. Global Change Research Program, Washington, DC, pp. 129–156.

Biagini, P., 2009. Classification of TTV and related viruses (anelloviruses). Current Topics in Microbiology and Immunology 331, 21–33.

Breitbart, M., Delwart, E., Rosario, K., Segales, J., Varsani, A., ICTV Report Consortium, 2017. ICTV Virus Taxonomy Profile: *Circoviridae*. Journal of General Virology 98, 1997–1998.

Burnard, D., Shao, R., 2019. Mitochondrial genome analysis reveals intraspecific variation within Australian hard tick species. Ticks and Tick-borne Diseases 10, 677–681.

Burnard, D., Weaver, H., Gillett, A., Loader, J., Flanagan, C., Polkinghorne, A., 2017. Novel Chlamydiales genotypes identified in ticks from Australian wildlife. Parasites & Vectors 10, 46.

Camacho, C., Coulouris, G., Avagyan, V., Ma, N., Papadopoulos, J., Bealer, K., Madden, T.L., 2009. BLAST+: architecture and applications. BMC Bioinformatics 10, 421.

Capella-Gutierrez, S., Silla-Martinez, J.M., Gabaldon, T., 2009. trimAl: a tool for automated alignment trimming in large-scale phylogenetic analyses. Bioinformatics 25, 1972–1973.

Cebria-Mendoza, M., Beamud, B., Andreu-Moreno, I., Arbona, C., Larrea, L., Diaz, W., Sanjuan, R., Cuevas, J.M., 2023. Human Anelloviruses: Influence of Demographic Factors, Recombination, and Worldwide Diversity. Microbiology Spectrum 11, e0492822.

Chandra, S., Harvey, E., Emery, D., Holmes, E.C., Slapeta, J., 2021. Unbiased Characterization of the Microbiome and Virome of Questing Ticks. Frontiers in Microbiology 12, 627327.

Clausen, P., Aarestrup, F.M., Lund, O., 2018. Rapid and precise alignment of raw reads against redundant databases with KMA. BMC Bioinformatics 19, 307.

Dehhaghi, M., Kazemi Shariat Panahi, H., Holmes, E.C., Hudson, B.J., Schloeffel, R., Guillemin, G.J., 2019. Human Tick-Borne Diseases in Australia. Frontiers in Cellular and Infection Microbiology 9, 3.

Diop, A., El Karkouri, K., Raoult, D., Fournier, P.E., 2020. Genome sequence-based criteria for demarcation and definition of species in the genus *Rickettsia*. International Journal of Systematic and Evolutionary Microbiology 70, 4249–4256.

Dong, X., El Karkouri, K., Robert, C., Raoult, D., Fournier, P.E., 2012. Genome sequence of *Rickettsia australis*, the agent of Queensland tick typhus. Journal of Bacteriology 194, 5129.

Elith, J., Leathwick, J.R., 2009. Species Distribution Models: Ecological Explanation and Prediction Across Space and Time. Annual Review of Ecology, Evolution, and Systematics 40, 677–697.

Estrada-Peña, A., Horak, I.G., Petney, T., 2008. Climate changes and suitability for the ticks *Amblyomma hebraeum* and *Amblyomma variegatum* (Ixodidae) in Zimbabwe (1974–1999). Veterinary Parasitology 151, 256–267.

Graham, R.M.A., Donohue, S., McMahon, J., Jennison, A.V., 2017. Detection of Spotted Fever Group *Rickettsia* DNA by Deep Sequencing. Emerging Infectious Diseases 23, 1911–1913.

Graves, S., Stenos, J., 2009. Rickettsioses in Australia. Annals of the New York Academy of Sciences 1166, 151–155.

Graves, S.R., Stenos, J., 2017. Tick-borne infectious diseases in Australia. Medical Journal of Australia 206, 320–324.

Greay, T.L., Oskam, C.L., Gofton, A.W., Rees, R.L., Ryan, U.M., Irwin, P.J., 2016. A survey of ticks (Acari: Ixodidae) of companion animals in Australia. Parasites & Vectors 9, 207.

Hahn, M.B., Jarnevich, C.S., Monaghan, A.J., Eisen, R.J., 2016. Modeling the geographic distribution of *Ixodes scapularis* and *Ixodes pacificus* (Acari: Ixodidae) in the contiguous United States. Journal of Medical Entomology 53, 1176–1191.

Harding, E.F., Russo, A.G., Yan, G.J.H., Waters, P.D., White, P.A., 2021. Ancient viral integrations in marsupials: a potential antiviral defence. Virus Evolution 7, veab076.

Harvey, E., Rose, K., Eden, J.S., Lo, N., Abeyasuriya, T., Shi, M., Doggett, S.L., Holmes, E.C., 2019. Extensive Diversity of RNA Viruses in Australian Ticks. Journal of Virology 93, e01358–18.

Hijmans, R.J., Phillips, S., Barbosa, M., Brunsdon, C., Rowlingson, B., 2024. predicts: spatial prediction tools (version 0.1-17).

Hirzel, A.H., Le Lay, G., Helfer, V., Randin, C., Guisan, A., 2006. Evaluating the ability of habitat suitability models to predict species presences. Ecological Modelling 199, 142–152.

Izzard, L., Graves, S., Cox, E., Fenwick, S., Unsworth, N., Stenos, J., 2009. Novel rickettsia in ticks, Tasmania, Australia. Emerging Infectious Diseases 15, 1654–1656.

Jansen van Vuren, P., Parry, R., Khromykh, A.A., Paweska, J.T., 2021. A 1958 Isolate of Kedougou Virus (KEDV) from Ndumu, South Africa, Expands the Geographic and Temporal Range of KEDV in Africa. Viruses 13, 685.

Jeffrey, S.J., Carter, J.O., Moodie, K.B., Beswick, A.R., 2001. Using spatial interpolation to construct a comprehensive archive of Australian climate data. Environmental Modelling & Software 16, 309–330.

Kaczorowska, J., van der Hoek, L., 2020. Human anelloviruses: diverse, omnipresent and commensal members of the virome. FEMS Microbiology Reviews 44, 305–313.

Kalyaanamoorthy, S., Minh, B.Q., Wong, T.K.F., von Haeseler, A., Jermiin, L.S., 2017. ModelFinder: fast model selection for accurate phylogenetic estimates. Nature Methods 14, 587–589.

Kass, J.M., Muscarella, R., Galante, P.J., Bohl, C.L., Pinilla-Buitrago, G.E., Boria, R.A., Soley-Guardia, M., Anderson, R.P., 2021. ENMeval 2.0: Redesigned for customizable and reproducible modeling of species’ niches and distributions. Methods in Ecology and Evolution 12, 1602–1608.

Katoh, K., Rozewicki, J., Yamada, K.D., 2019. MAFFT online service: multiple sequence alignment, interactive sequence choice and visualization. Briefings in Bioinformatics 20, 1160–1166.

Lee, I., Ouk Kim, Y., Park, S.C., Chun, J., 2016. OrthoANI: An improved algorithm and software for calculating average nucleotide identity. International Journal of Systematic and Evolutionary Microbiology 66, 1100–1103.

Leta, S., De Clercq, E.M., Madder, M., 2013. High-resolution predictive mapping for *Rhipicephalus appendiculatus* (Acari: Ixodidae) in the Horn of Africa. Experimental and Applied Acarology 60, 531–542.

Li, D., Liu, C.M., Luo, R., Sadakane, K., Lam, T.W., 2015. MEGAHIT: an ultra-fast single-node solution for large and complex metagenomics assembly via succinct de Bruijn graph. Bioinformatics 31, 1674–1676.

Low, B.W., Zeng, Y., Tan, H.H., Yeo, D.C.J., 2021. Predictor complexity and feature selection affect Maxent model transferability: evidence from global freshwater invasive species. Diversity and Distributions 27, 497–511.

Ma, R., Li, C., Gao, A., Jiang, N., Li, J., Hu, W., Feng, X., 2024. Tick species diversity and potential distribution alternation of dominant ticks under different climate scenarios in Xinjiang, China. PLoS Neglected Tropical Diseases 18, e0012108.

Ma, R., Li, C., Tian, H., Zhang, Y., Feng, X., Li, J., Hu, W., 2023. The current distribution of tick species in Inner Mongolia and inferring potential suitability areas for dominant tick species based on the MaxEnt model. Parasites & Vectors 16, 286.

Marcelino, V.R., Clausen, P., Buchmann, J.P., Wille, M., Iredell, J.R., Meyer, W., Lund, O., Sorrell, T.C., Holmes, E.C., 2020. CCMetagen: comprehensive and accurate identification of eukaryotes and prokaryotes in metagenomic data. Genome Biology 21, 103.

Marchler-Bauer, A., Lu, S., Anderson, J.B., Chitsaz, F., Derbyshire, M.K., DeWeese-Scott, C., Fong, J.H., Geer, L.Y., Geer, R.C., Gonzales, N.R., Gwadz, M., Hurwitz, D.I., Jackson, J.D., Ke, Z., Lanczycki, C.J., Lu, F., Marchler, G.H., Mullokandov, M., Omelchenko, M.V., Robertson, C.L., Song, J.S., Thanki, N., Yamashita, R.A., Zhang, D., Zhang, N., Zheng, C., Bryant, S.H., 2011. CDD: a Conserved Domain Database for the functional annotation of proteins. Nucleic Acids Research 39, D225–229.

McInnes, L.M., Hanger, J., Simmons, G., Reid, S.A., Ryan, U.M., 2011. Novel trypanosome *Trypanosoma gilletti* sp. (Euglenozoa: Trypanosomatidae) and the extension of the host range of *Trypanosoma copemani* to include the koala (*Phascolarctos cinereus*). Parasitology 138, 59–70.

Ng, T.F., Willner, D.L., Lim, Y.W., Schmieder, R., Chau, B., Nilsson, C., Anthony, S., Ruan, Y., Rohwer, F., Breitbart, M., 2011. Broad surveys of DNA viral diversity obtained through viral metagenomics of mosquitoes. PLoS One 6, e20579.

O’Brien, C.A., Hall-Mendelin, S., Hobson-Peters, J., Deliyannis, G., Allen, A., Lew-Tabor, A., Rodriguez-Valle, M., Barker, D., Barker, S.C., Hall, R.A., 2018. Discovery of a novel iflavirus sequence in the eastern paralysis tick *Ixodes holocyclus*. Archives of Virology 163, 2451–2457.

Okamoto, H., Nishizawa, T., Takahashi, M., Tawara, A., Peng, Y., Kishimoto, J., Wang, Y., 2001. Genomic and evolutionary characterization of TT virus (TTV) in tupaias and comparison with species-specific TTVs in humans and non-human primates. Journal of General Virology 82, 2041–2050.

Parks, D.H., Imelfort, M., Skennerton, C.T., Hugenholtz, P., Tyson, G.W., 2015. CheckM: assessing the quality of microbial genomes recovered from isolates, single cells, and metagenomes. Genome Research 25, 1043–1055.

Parola, P., Paddock, C.D., Socolovschi, C., Labruna, M.B., Mediannikov, O., Kernif, T., Abdad, M.Y., Stenos, J., Bitam, I., Fournier, P.E., Raoult, D., 2013. Update on tick-borne rickettsioses around the world: a geographic approach. Clinical Microbiology Reviews 26, 657–702.

Parry, R., James, M.E., Asgari, S., 2021. Uncovering the Worldwide Diversity and Evolution of the Virome of the Mosquitoes *Aedes aegypti* and *Aedes albopictus*. Microorganisms 9, 1236.

Pascoe, E.L., Nava, S., Labruna, M.B., Paddock, C.D., Levin, M.L., Marcantonio, M., Foley, J.E., 2022. Predicting the northward expansion of tropical lineage *Rhipicephalus sanguineus sensu lato* ticks in the United States and its implications for medical and veterinary health. PLOS ONE 17, e0271683.

Perret, J.L., Guigoz, E., Rais, O., Gern, L., 2000. Influence of saturation deficit and temperature on *Ixodes ricinus* tick questing activity in a Lyme borreliosis-endemic area (Switzerland). Parasitology Research 86, 554–557.

Phillips, S., Quigley, B.L., Aziz, A., Bergen, W., Booth, R., Pyne, M., Timms, P., 2019. Antibiotic treatment of Chlamydia-induced cystitis in the koala is linked to expression of key inflammatory genes in reactive oxygen pathways. PLoS One 14, e0221109.

Phillips, S.J., Anderson, R.P., Dudík, M., Schapire, R.E., Blair, M.E., 2017. Opening the black box: an open-source release of Maxent. Ecography 40, 887–893.

Raghavan, R.K., Barker, S.C., Cobos, M.E., Barker, D., Teo, E.J.M., Foley, D.H., Nakao, R., Lawrence, K., Heath, A.C.G., Peterson, A.T., 2019. Potential spatial distribution of the newly introduced long-horned tick, *Haemaphysalis longicornis* in North America. Scientific Reports 9, 498.

Randolph, S.E., Storey, K., 1999. Impact of microclimate on immature tick-rodent host interactions (Acari: Ixodidae): implications for parasite transmission. Journal of Medical Entomology 36, 741–748.

Robinson, S.J., Neitzel, D.F., Moen, R.A., Craft, M.E., Hamilton, K.E., Johnson, L.B., Mulla, D.J., Munderloh, U.G., Redig, P.T., Smith, K.E., Turner, C.L., Umber, J.K., Pelican, K.M., 2015. Disease risk in a dynamic environment: the spread of tick-borne pathogens in Minnesota, USA. Ecohealth 12, 152–163.

Rochlin, I., Egizi, A., Narvaez, Z., Bonilla, D.L., Gallagher, M., Williams, G.M., Rainey, T., Price, D.C., Fonseca, D.M., 2023. Microhabitat modeling of the invasive Asian longhorned tick (*Haemaphysalis longicornis*) in New Jersey, USA. Ticks and Tick-borne Diseases 14, 102126.

Segales, J., 2012. Porcine circovirus type 2 (PCV2) infections: clinical signs, pathology and laboratory diagnosis. Virus Research 164, 10–19.

Sentausa, E., Abdad, M.Y., Robert, C., Stenos, J., Raoult, D., Fournier, P.E., 2013. Genome Sequence of *Rickettsia gravesii*, Isolated from Western Australian Ticks. Genome Announcements 1, e00097–13.

Simao, F.A., Waterhouse, R.M., Ioannidis, P., Kriventseva, E.V., Zdobnov, E.M., 2015. BUSCO: assessing genome assembly and annotation completeness with single-copy orthologs. Bioinformatics 31, 3210–3212.

Spandole, S., Cimponeriu, D., Berca, L.M., Mihaescu, G., 2015. Human anelloviruses: an update of molecular, epidemiological and clinical aspects. Archives of Virology 160, 893–908.

Stewart, A., Armstrong, M., Graves, S., Hajkowicz, K., 2017a. Epidemiology and Characteristics of *Rickettsia australis* (Queensland Tick Typhus) Infection in Hospitalized Patients in North Brisbane, Australia. Tropical Medicine and Infectious Disease 2, 35.

Stewart, A., Armstrong, M., Graves, S., Hajkowicz, K., 2017b. *Rickettsia australis* and Queensland Tick Typhus: A Rickettsial Spotted Fever Group Infection in Australia. American Journal of Tropical Medicine and Hygiene 97, 24–29.

Stewart, A.G., Stewart, A.G.A., 2021. An Update on the Laboratory Diagnosis of *Rickettsia* spp. Infection. Pathogens 10, 1280.

Stewart, A.G.A., Smith, S., Binotto, E., McBride, W.J.H., Hanson, J., 2019. The epidemiology and clinical features of rickettsial diseases in North Queensland, Australia: Implications for patient identification and management. PLoS Neglected Tropical Diseases 13, e0007583.

Tatusova, T., DiCuccio, M., Badretdin, A., Chetvernin, V., Nawrocki, E.P., Zaslavsky, L., Lomsadze, A., Pruitt, K.D., Borodovsky, M., Ostell, J., 2016. NCBI prokaryotic genome annotation pipeline. Nucleic Acids Research 44, 6614–6624.

Taylor-Brown, A., Bachmann, N.L., Borel, N., Polkinghorne, A., 2016. Culture-independent genomic characterisation of Candidatus *Chlamydia sanzinia*, a novel uncultivated bacterium infecting snakes. BMC Genomics 17, 710.

Teo, E.J.M., Apanaskevich, D.A., Barker, S.C., Nakao, R., 2024a. *Dermacentor* (*Indocentor*) *auratus* Supino 1897: Potential geographic range, and medical and veterinary significance. Acta Tropica 254, 107197.

Teo, E.J.M., Hailu, S., Kelava, S., Zalucki, M.P., Furlong, M.J., Nakao, R., Barker, D., Barker, S.C., 2021a. Climatic requirements of the southern paralysis tick, *Ixodes cornuatus*, with a consideration of its host, *Vombatus ursinus*, and the possible geographic range of the tick up to 2090. Ticks and Tick-borne Diseases 12, 101758.

Teo, E.J.M., Russell, H., Lambert, T., Webster, R., Yappa, A., McDonagh, P., Harper, G., Barker, D., Barker, S.C., 2023. The weather determines the number of cases of tick paralysis in dogs and cats in eastern Australia, caused by *Ixodes holocyclus*, the eastern paralysis tick. Australian Veterinary Journal 101, 513–523.

Teo, E.J.M., Russell, H., Lambert, T., Webster, R., Yappa, A., McDonagh, P., Harper, G., Barker, D., Nakao, R., Barker, S.C., 2024b. The weather determined how ‘hot’ the tick paralysis season was in eastern Australia: 2018–2024. Veterinary Parasitology 331, 110252.

Teo, E.J.M., Vial, M.N., Hailu, S., Kelava, S., Zalucki, M.P., Furlong, M.J., Barker, D., Barker, S.C., 2021b. Climatic requirements of the eastern paralysis tick, *Ixodes holocyclus*, with a consideration of its possible geographic range up to 2090. International Journal of Parasitology 51, 241–249.

Todd, D., Weston, J., Ball, N.W., Borghmans, B.J., Smyth, J.A., Gelmini, L., Lavazza, A., 2001. Nucleotide sequence-based identification of a novel circovirus of canaries. Avian Pathology 30, 321–325.

Valavi, R., Elith, J., Lahoz-Monfort, J.J., Guillera-Arroita, G., 2019. blockCV: An r package for generating spatially or environmentally separated folds for k-fold cross-validation of species distribution models. Methods in Ecology and Evolution 10, 225–232.

Varsani, A., Opriessnig, T., Celer, V., Maggi, F., Okamoto, H., Blomstrom, A.L., Cadar, D., Harrach, B., Biagini, P., Kraberger, S., 2021. Taxonomic update for mammalian anelloviruses (family Anelloviridae). Archives of Virology 166, 2943–2953.

Vilcins, I.M., Old, J.M., Deane, E., 2009. Detection of a Hepatozoon and spotted fever group *Rickettsia* species in the common marsupial tick (*Ixodes tasmani*) collected from wild Tasmanian devils (*Sarcophilus harrisii*), Tasmania. Veterinary Parasitology 162, 23–31.

Vilcins, I.M., Old, J.M., Deane, E.M., 2008. Detection of a spotted fever group *Rickettsia* in the tick *Ixodes tasmani* collected from koalas in Port Macquarie, Australia. Journal of Medical Entomology 45, 745–750.

Waits, K., Edwards, M.J., Cobb, I.N., Fontenele, R.S., Varsani, A., 2018. Identification of an anellovirus and genomoviruses in ixodid ticks. Virus Genes 54, 155–159.

Wang, B., Sun, L.D., Liu, H.H., Wang, Z.D., Zhao, Y.K., Wang, W., Liu, Q., 2018. Molecular detection of novel circoviruses in ticks in northeastern China. Ticks and Tick-borne Diseases 9, 836–839.

Westgate, M., Kellie, D., Stevenson, M., Newman, P., 2025. galah: biodiversity data from the GBIF node network. R package version 2.1.1.

Willis, G., Lodo, K., McGregor, A., Howes, F., Williams, S., Veitch, M., 2019. New and old hotspots for rickettsial spotted fever acquired in Tasmania, 2012-2017. Australian and New Zealand Journal of Public Health 43, 389–394.

Wu, Y.W., Simmons, B.A., Singer, S.W., 2016. MaxBin 2.0: an automated binning algorithm to recover genomes from multiple metagenomic datasets. Bioinformatics 32, 605–607.

Xin, D., El Karkouri, K., Robert, C., Raoult, D., Fournier, P.E., 2012. Genomic comparison of *Rickettsia honei* strain RBT and other *Rickettsia* Species. Journal of Bacteriology 194, 4145.

Xu, L., Guo, Y., Yang, L., Li, Z., Kang, M., Han, X., Chen, C., He, S., Hu, X., He, Y., Wang, Y., Li, Z., Chen, J., Geng, P., Chen, Q., Jiang, S., Ma, J., Zhang, X., Tai, X., Li, Y., 2024. Projected distribution and dispersal patterns of prevalent ticks and tick-borne pathogens in the Sanjiangyuan area of Qinghai province, China, under intense climatic conditions. Frontiers in Environmental Science 12, 1309290.

Zemtsova, G.E., Apanaskevich, D.A., Reeves, W.K., Hahn, M., Snellgrove, A., Levin, M.L., 2016. Phylogeography of *Rhipicephalus sanguineus sensu lato* and its relationships with climatic factors. Experimental and Applied Acarology 69, 191–203.

Zhao, M., Yue, C., Yang, Z., Li, Y., Zhang, D., Zhang, J., Yang, S., Shen, Q., Su, X., Qi, D., Ma, R., Xiao, Y., Hou, R., Yan, X., Li, L., Zhou, Y., Liu, J., Wang, X., Wu, W., Zhang, W., Shan, T., Liu, S., 2022. Viral metagenomics unveiled extensive communications of viruses within giant pandas and their associated organisms in the same ecosystem. Science of the Total Environment 820, 153317.

Zuker, M., 2003. Mfold web server for nucleic acid folding and hybridization prediction. Nucleic Acids Research 31, 3406–3415.

